# The *Gossypium longicalyx* genome as a resource for cotton breeding and evolution

**DOI:** 10.1101/2020.01.08.898908

**Authors:** Corrinne E. Grover, Mengqiao Pan, Daojun Yuan, Mark A. Arick, Guanjing Hu, Logan Brase, David M. Stelly, Zefu Lu, Robert J. Schmitz, Daniel G. Peterson, Jonathan F. Wendel, Joshua A. Udall

## Abstract

Cotton is an important crop that has made significant gains in production over the last century. Emerging pests such as the reniform nematode have threatened cotton production. The rare African diploid species *Gossypium longicalyx* is a wild species that has been used as an important source of reniform nematode immunity. While mapping and breeding efforts have made some strides in transferring this immunity to the cultivated polyploid species, the complexities of interploidal transfer combined with substantial linkage drag have inhibited progress in this area. Moreover, this species shares its most recent common ancestor with the cultivated A-genome diploid cottons, thereby providing insight into the evolution of long, spinnable fiber. Here we report a newly generated *de novo* genome assembly of *G. longicalyx*. This high-quality genome leveraged a combination of PacBio long-read technology, Hi-C chromatin conformation capture, and BioNano optical mapping to achieve a chromosome level assembly. The utility of the *G. longicalyx* genome for understanding reniform immunity and fiber evolution is discussed.

## Introduction

Cotton (genus *Gossypium*) is an important crop which provides the largest natural source of fiber. Colloquially, the term cotton refers to one of four domesticated species, primarily the tetraploid *G. hirsutum*, which is responsible for over 98% of cotton production worldwide (Kranthi 2018). *Gossypium* contains over 50 additional wild species related to the domesticated cottons that serve as potential sources of disease and pest resistance. Among these, *Gossypium longicalyx* J.B. Hutch. & B.J.S. Lee is the only representative of the diploid “F-genome” (Wendel and Grover 2015) and the only species with immunity to reniform nematode infection (Yik and Birchfield 1984). Discovered only 60 years ago (Hutchinson and B. J. S. Lee 1958), it is both cytogenetically differentiated from members of the other genome groups (Phillips 1966) and morphologically isolated (Fryxell 1971, 1992). Importantly, *G. longicalyx* is sister to the A-genome cottons (Wendel and Albert 1992; Wendel and Grover 2015; Chen *et al.* 2016), i.e., *G. arboreum* and *G. herbaceum*, the only diploids with long, spinnable fiber.

Interest in the genome of *G. longicalyx* is two-fold. First, broad-scale screening of the cotton germplasm collection indicates that domesticated cotton lacks appreciable natural resistance to reniform nematode (Birchfield *et al.* 1963; Yik and Birchfield 1984), and while several other species exhibit degrees of resistance, only *G. longicalyx* exhibits immunity to infection (Yik and Birchfield 1984). This is significant as reniform nematode has emerged as a major source of cotton crop damage, reducing cotton production by over 205 million bales per year (Lawrence *et al.* 2015) and accounting for ~11% of the loss attributable to pests (Khanal *et al.* 2018). Current reniform resistant lines are derived from complex breeding schemes which are required to introgress reniform immunity from the diploid *G. longicalyx* into polyploid *G. hirsutum* (Bell and Robinson 2004; Dighe *et al.* 2009; Khanal *et al.* 2018); however, undesirable traits have accompanied this introgression (Nichols *et al.* 2010) extreme stunting of seedlings and plants exposed to dense nematode populations, prohibiting commercial deployment (Zheng *et al.* 2016).

The genome of *G. longicalyx* is also valuable because it is phylogenetically sister to the only diploid clade with spinnable fiber (Wendel and Albert, 1992; Wendel and Grover, 2015; Chen et al., 2016), the A-genome species, which contributed the maternal ancestor to polyploid cotton. Consequently, there has been interest in this species as the ancestor to spinnable fiber (Hovav *et al.* 2008; Paterson *et al.* 2012), although progress has been limited due to lack of genomic resources in *G. longicalyx*. Comparisons between the *G. longicalyx* genome and other cotton genomes, including the domesticated diploids (Du *et al.* 2018), may provide clues into the evolutionary origin of “long” fiber.

Here we describe a high-quality, *de novo* genome sequence for *G. longicalyx*, a valuable resource for understanding nematode immunity in cotton and possibly other species. This genome also provides a foundation to understand the evolutionary origin of spinnable fiber in *Gossypium*.

## Methods & Materials

### Plant material and sequencing methods

Leaf tissue of mature *G. longicalyx* (F1-1) was collected from a Brigham Young University (BYU) greenhouse. DNA was extracted using CTAB techniques (Kidwell and Osborn 1992), and the amount recovered was measured via Qubit Fluorometer (ThermoFisher, Inc.). The sequencing library was constructed by the BYU DNA Sequencing Center (DNASC) using only fragments >18 kb, which were size selected on the BluePippen (Sage Science, LLC) and verified in size using a Fragment Analyzer (Advanced Analytical Technologies, Inc). Twenty-six PacBio cells were sequenced from a single library on the Pacific Biosciences Sequel system. Resulting reads were assembled using Canu V1.6 using default parameters (Koren *et al.* 2017) to create a sequence assembly called Longicalyx_V1.0, composed of 229 large contigs (Figure 1).

**Figure 1.**
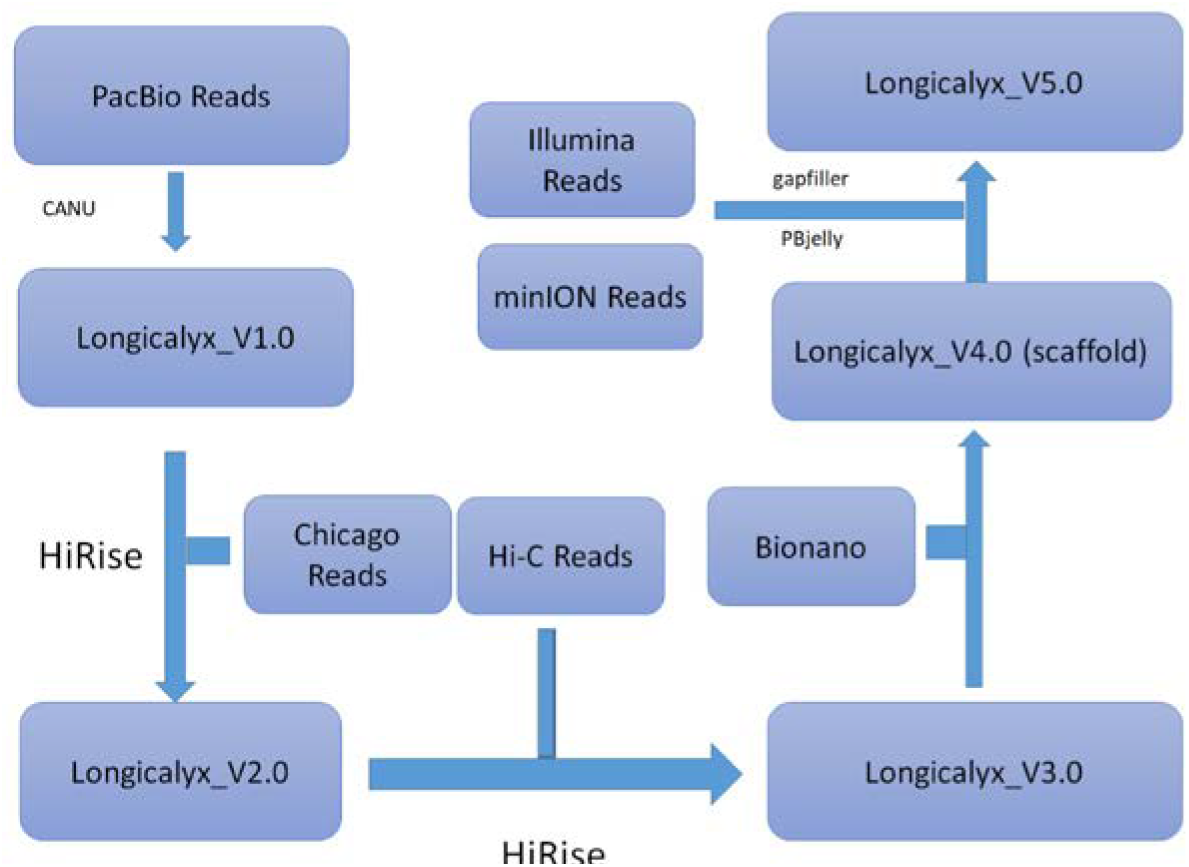
Chicago Highrise reads (Dovetail Genomics) provide DNA-DNA proximity information used to improve the Canu sequence assembly (Longicalyx_V2.0; statistics not calculated), as previously demonstrated for *de novo* human and alligator genomes (Putnam *et al.* 2016). Simultaneously, HiC libraries were constructed from *G. longicalyx* leaf tissue at PhaseGenomics LLC. A second round of HighRise was used to include the HiC data for additional genome scaffolding (Koch 2016; Putnam *et al.* 2016), reducing the contig number to 135 (Longicalyx_V3.0).

High-molecular weight DNA was extracted from young *G. longicalyx* leaves and subsequently purified, nicked, labeled, and repaired according to Bionano Plant protocol and standard operating procedures for the Irys platform. BssSI was used in conjunction with the IrysSolve pipeline to assemble an optical map on the BYU Fulton SuperComputing cluster. The resulting optical map was aligned to the assembly named Lonigcalyx_V3.0 using an *in silico* labeled reference sequence. Bionano maps linked large contigs present in this assembly, producing 17 large scaffolds (Lonigcalyx_V4.0).

Minion sequencing libraries were created and sequenced following the standard protocol from Oxford Nanopore. Scaffolds from Lonigcalyx_V4.0 were polished (Supplemental File 1) with existing Illumina (SRR1174179 and SRR1174182 from the NCBI Short Read Archive) and the newly generated Minion data for *G. longicalyx* using both PBjelly (English *et al.* 2012) and GapFiller (Boetzer and Pirovano 2012) to produce the final assembly, Lonigcalyx_V5.0.

### Repeat and gene annotation

Repeats were identified using two methods. The first is a homology-based approach, *i.e.*, a combination of RepeatMasker (Smit *et al.* 2015) and “One code to find them all” (Bailly-Bechet *et al.* 2014), whereas the second method (i.e., RepeatExplorer; (Novák *et al.* 2010) clusters reads based on sequence similarity and automatically annotates the most abundant cluster using RepeatMasker. Each RepeatMasker run used a custom library, which combines Repbase 23.04 repeats (Bao *et al.* 2015) with cotton-specific repeats. Default parameters were run, except the run was “sensitive” and was set to mask only TEs (no low-complexity). Parameters are available https://github.com/Wendellab/longicalyx. “One code to find them all” was used to aggregate multiple hits from the first method (RepeatMasker) into TE models using default parameters. The resulting output was aggregated and summarized in R/3.6.0 (R Core Team 2017) using *dplyr */0.8.1 (Wickham *et al.* 2015). Cluster results were obtained from (Grover *et al.* 2019) and https://github.com/IGBB/D_Cottons_USDA, and these were parsed in R/3.6.0 (R Core Team 2017). All code is available at https://github.com/Wendellab/longicalyx.

RNA-Seq libraries were generated from *G. longicalyx* leaf (CL), floral (FF), and stem tissues (FS) to improve genome annotation. RNA-seq libraries were independently constructed by BGI Americas (Davis, CA) using Illumina TruSeq reagents and subsequently sequenced (single-end, 50 bp). The newly sequenced *G. longicalyx* RNA-seq was combined with existing RNA-seq from *G. longicalyx* (SRR1174179) as well as two closely related species, i.e., *G. herbaceum* (developing fibers and seed; PRJNA595350 and SRR959585, respectively) and *G. arboreum* (5 seed libraries and 1 seedling; SRR617075, SRR617073, SRR617068, SRR617067, SRR959590, and SRR959508). RNA-seq libraries were mapped to the hard-masked *G. longicalyx* genome using hisat2 [v2.1.0] (Kim *et al.* 2015). BRAKER2 [v2.1.2] (Hoff *et al.* 2019) was used in conjunction with GeneMark [v4.36] (Borodovsky and Lomsadze 2011) generated annotations to train Augustus [v3.3.2] (Stanke *et al.* 2006). Mikado [v1.2.4] (Venturini *et al.* 2018) was used to produce high quality RNA-seq based gene predictions by combining the RNA-seq assemblies produced by StringTie [v1.3.6] (Pertea *et al.* 2015) and Cufflinks [v2.2.1] (Ghosh and Chan 2016) with a reference-guided assembly from Trinity [v2.8.5] (Grabherr *et al.* 2011) and a splice junction analysis from Portcullis [v1.2.2] (Mapleson *et al.* 2018). The Trinity assembly was formatted using GMAP [v2019-05-12] (Wu and Watanabe 2005). MAKER2 [v2.31.10] (Holt and Yandell 2011; Campbell *et al.* 2014) was used to integrate gene predictions from (1) BRAKER2 trained Augustus, (2) GeneMark, and (3) Mikado, also using evidence from all *Gossypium* ESTs available from NCBI (nucleotide database filtered on “txid3633” and “is_est”) and a database composed of all curated proteins in Uniprot Swissprot [v2019_07] (UniProt Consortium 2008) combined with the annotated proteins from the *G. hirsutum* (https://www.cottongen.org/species/Gossypium_hirsutum/jgi-AD1_genome_v1.1) and *G. raimondii* (Paterson *et al.* 2012) genomes. Maker scored each gene model using the annotation edit distance (AED - (Eilbeck *et al.* 2009; Holt and Yandell 2011; Yandell and Ence 2012) metric based on EST and protein evidence provided. Gene models with an AED greater than 0.47 were removed from further analyses, and the remaining gene models were functionally annotated using InterProScan [v5.35-74.0] (Jones *et al.* 2014) and BlastP [v2.9.0+] (Camacho *et al.* 2009) searches against the Uniprot SwissProt database. Orthologs between the *G. longicalyx* annotations and the existing annotations for *G. arboreum (Du et al. 2018)*, *G. raimondii* (Paterson *et al.* 2012)*, G. hirsutum* (Hu *et al.* 2019), and *G. barbadense* (Hu *et al.* 2019) were predicted by OrthoFinder using default settings (Emms and Kelly 2015, 2019). All genomes are hosted through CottonGen (https://www.cottongen.org; (Yu *et al.* 2014)) and running parameters are available from https://github.com/Wendellab/longicalyx.

### ATAC-seq and data analysis

ATAC-seq was performed as described previously (Lu *et al.* 2017). For each replicate, approximately 200 mg freshly collected leaves or flash frozen leaves were immediately chopped with a razor blade in ~ 1 ml of pre-chilled lysis buffer (15 mM Tris-HCl pH 7.5, 20 mM NaCl, 80 mM KCl, 0.5 mM spermine, 5 mM 2-mercaptoethanol, 0.2% Triton X-100). The chopped slurry was filtered twice through miracloth and once through a 40 μm filter. The crude nuclei were stained with DAPI and loaded into a flow cytometer (Beckman Coulter MoFlo XDP). Nuclei were purified by flow sorting and washed in accordance with Lu et al (Lu *et al.* 2017). The sorted nuclei were incubated with 2 μl Tn5 transposomes in 40 μl of tagmentation buffer (10 mM TAPS-NaOH ph 8.0, 5 mM MgCl_2_) at 37°C for 30 minutes without rotation. The integration products were purified using a Qiagen MinElute PCR Purification Kit or NEB Monarch™ DNA Cleanup Kit and then amplified using Phusion DNA polymerase for 10-13 cycles. PCR cycles were determined as described previously (Buenrostro *et al.* 2013). Amplified libraries were purified with AMPure beads to remove primers. ATAC-seq libraries were sequenced in paired-end 35 bp at the University of Georgia Genomics & Bioinformatics Core using an Illumina NextSeq 500 instrument.

Reads were adapter and quality trimmed, and then filtered using “Trim Galore” [v0.4.5] (Krueger 2015). Clean reads were subsequently aligned to the Lonigcalyx_V5.0 assembly using Bowtie2 [v2.3.4] (Langmead and Salzberg 2012) with the parameters “--no-mixed --no-discordant --no-unal --dovetail”. Duplicate reads were removed using Picard [v2.17.0] with default parameters (http://broadinstitute.github.io/picard/). Only uniquely mapped read pairs with a quality score of at least 20 were kept for peak calling. Phantompeakqualtools [v1.14] (Landt *et al.* 2012) was used to calculate the strand cross-correlation, and deepTools [v2.5.2] (Ramírez *et al.* 2016) was used to calculate correlation between replicates. The peak calling tool from HOMER [v4.10] (Heinz *et al.* 2010), i.e., *findpeaks*, was run in “region” mode and with the minimal distance between peaks set to 150 bp. MACS2 [v2.1.1] (Zhang *et al.* 2008) *callpeak*, a second peak-calling algorithm, was run with the parameter “-f BAMPE” to analyze only properly paired alignments, and putative peaks were filtered using default settings and false discovery rate (FDR) < 0.05. Due to the high level of mapping reproducibility (Pearson’s correlation *r* = 0.99 and Spearman correlation *r* = 0.77 by deepTools), peaks were combined and merged between replicates for each tool using BEDTools [v2.27.1] (Quinlan 2014). BEDTools was also used to intersect HOMER peaks and MACS2 peaks to only retain peak regions identified by both tools as accessible chromatin regions (ACRs) for subsequent analyses.

ACRs were annotated in relation to the nearest annotated genes in the R environment [v3.5.0] as genic (gACRs; overlapping a gene), proximal (pACRs; within 2 Kb of a gene) or distal (dACRs; >2 Kb from a gene). Using R package ChIPseeker [v1.18.0] (Yu *et al.* 2015), the distribution of ACRs was calculated around transcription start sites (TSS) and transcription termination sites (TTS), and peak distribution was visualized with aggregated profiles and heatmaps. To compare GC contents between ACRs and non-accessible genomic region, the BEDTools *shuffle* command was used to generate the distal (by excluding genic and 2 Kb flanking regions) and genic/proximal control regions (by including genic and 2 Kb flanking regions), and the *nuc* command was used to calculate GC content for each ACR and permuted control regions.

### Identification of the Ren^Lon^ region in *G. longicalyx*

Previous research (Dighe *et al.* 2009; Zheng *et al.* 2016) identified a marker (BNL1231) that consistently cosegrates with resistance and that is flanked by the SNP markers Gl_168758 and Gl_072641, which are all located in the region of *G. longicalyx* chromosome 11 referred to as “Ren^Lon^”. These three markers were used as queries of gmap (Wu and Watanabe 2005) against the assembled genome to identify the genomic regions associated with each. The coordinates identified by gmap were placed in a bed file; this file was used in conjunction with the *G. longicalyx* annotation and BEDtools intersect (Quinlan 2014) to identify predicted *G. longicalyx* genes contained within Ren^Lon^. Samtools faidx (Li *et al.* 2009) was used to extract the 52 identified genes from the annotation file, which were functionally annotated using blast2go (blast2go basics; biobam) and including blastx (Altschul *et al.* 1990), gene ontology (The Gene Ontology Consortium 2019), and InterPro (Jones *et al.* 2014). Orthogroups containing each of the 52 Ren^Lon^ genes were identified from the Orthofinder results (see above).

### Comparison between *G. arboreum* and *G. longicalyx* for fiber evolution

Whole-genome alignments were generated between *G. longicalyx* and either *G. arboreum*, *G. raimondii*, *G. turneri*, *G. hirsutum* (A-chromosomes), and *G. barbadense* (A-chromosomes) using Mummer (Marçais *et al.* 2018) and visualized using dotPlotly (https://github.com/tpoorten/dotPlotly) in R (version 3.6.0) (R Development Core Team and Others 2011). Divergence between *G. longicalyx* and *G. arboreum* or *G. raimondii* was calculated using orthogroups that contain a single *G. longicalyx* gene with a single *G. arboreum* and/or single *G. raimondii* gene. Pairwise alignments between *G. longicalyx* and *G. arboreum* or *G. raimondii* were generated using the *linsi* from MAFFT (Katoh and Standley 2013). Pairwise distances between *G. longicalyx* and *G. arboreum* and/or *G. raimondii* were calculated in R (version 3.6.0) using phangorn (Schliep 2011) and visualized using ggplot2 (Wickham 2016). To identify genes unique to species with spinnable fiber (i.e., *G. arboreum* and the polyploid species), we extracted any *G. arboreum* gene contained within orthogroups composed solely of *G. arboreum* or polyploid A-genome gene annotations, and subjected these to blast2go (as above). Syntenic conservation of genes contained within the Ren^Lon^ region, as compared to *G. arboreum*, was evaluated using GEvo as implemented in SynMap via COGE (Lyons and Freeling 2008; Haug-Baltzell *et al.* 2017).

### Data availability

The assembled genome sequence of *G. longicalyx* is available at NCBI SUB6483233 and CottonGen (https://www.cottongen.org/). The raw data for *G. longicalyx* are also available at NCBI PRJNA420071 for PacBio and Minion, and PRJNA420070 for RNA-Seq. Supplemental files are available from figshare.

## Results and Discussion

### Genome assembly and annotation

We report a *de novo* genome sequence for *G. longicalyx*. This genome was first assembled from ~144x coverage (raw) of PacBio reads, which alone produced an assembly consisting of 229 contigs with an N50 of 28.8MB (Table 1). The contigs were scaffolded using a combination of Chicago Highrise, Hi-C, and BioNano to produce a chromosome level assembly consisting of 17 contigs with an average length of 70.4 Mb (containing only 8.4kb of gap sequence). Thirteen of the chromosomes were assembled into single contigs. Exact placement of the three unscaffolded contigs (~100 kb) was not determined, but these remaining sequences were included in NCBI with the assembled chromosomes. The final genome assembly size was 1190.7 MB, representing over 90% of the estimated genome size (Hendrix and Stewart 2005).

**Table 1.** Statistics for assembly versions

| Method | *G. longicalyx* assemblies* |  |  |  |
| --- | --- | --- | --- | --- |
|  | Longicalyx_V1.0<br>PacBio/Canu | Longicalyx_V3.0<br>+Chicago HighRise+HiC | Longicalyx_V4.0<br>+BioNano | Longicalyx_V5.0<br>+Illumina+Minion |
| Coverage | 79.45 |  |  |  |
| Total Contig Number | 229 | 135 | 17 | 17 |
| Assembly Length** | 1196.17 Mb | 1196.19 Mb | 1190.66 Mb | 1190.67 Mb |
| Average Contig Length | 5.22 Mb | 8.86 Mb | 70.04 Mb | 70.04 Mb |
| Total Length of Ns | 0 | 18200 | 18000 | 8488 |
| N50 value is | 28.88 Mb | 95.88 Mb | 95.88 Mb | 95.88 Mb |
| N90 value is | 7.58 Mb | 76.48 Mb | 76.48 Mb | 76.29 Mb |

\* Statistics for Longicalyx\_V2.0 not calculated
\*\* Genome size for *G. longicalyx* is 1311 (Hendrix and Stewart, 2005)

BUSCO analysis of the completed genome (Waterhouse *et al.* 2017) recovered 95.8% complete BUSCOs (from the total of 2121 BUSCO groups searched; Table 2). Most BUSCOs (86.5%) were both complete and single copy, with only 9.3% BUSCOs complete and duplicated. Less than 5% of BUSCOs were either fragmented (1.4%) or missing (2.8%), indicating a general completeness of the genome. Genome contiguity was independently verified using the LTR Assembly Index (LAI) (Ou *et al.* 2018), which is a reference-free method to assess genome contiguity by evaluating the completeness of LTR-retrotransposon assembly within the genome. This method, applied to over 100 genomes in Phytozome, suggests that an LAI between 10 and 20 should be considered “reference-quality”; the *G. longicalyx* genome reported here received an LAI score of 10.74. Comparison of the *G. longicalyx* genome to published cotton genomes (Table 2) suggests that the quality of this assembly is similar or superior to other currently available cotton genomes.

**Table 2.** BUSCO and LAI scores for the *G. longicalyx* genome compared to existing cotton genomes.

|  | Complete BUSCO |  |  | Incomplete BUSCO |  | LAI score | Reference |
| --- | --- | --- | --- | --- | --- | --- | --- |
|  | Total | Single | Duplicated | Fragmented | Missing |  |  |
| *G. longicalyx* | 95.80% | 86.50% | 9.30% | 1.40% | 2.80% | 10.74 |  |
| *G. turneri* | 95.80% | 86.00% | 9.80% | 1.00% | 3.20% | 8.51 | (Udall *et al.* 2019) |
| *G. raimondii* (BYU) | 92.80% | 85.10% | 7.70% | 2.70% | 4.50% | 10.57 | (Udall *et al.* 2019) |
| *G. raimondii* (JGI) | 98.00% | 87.30% | 10.70% | 0.70% | 1.30% | 8.51 | (Paterson *et al.* 2012) |
| *G. arboreum* (CRI) | 94.70% | 85.20% | 9.50% | 1.00% | 4.30% | 12.59 | (Du *et al.* 2018) |
| *G. barbadense* 3-79 (HAU v2) | 96.30% | 12.20% | 84.10% | 0.80% | 2.90% | 10.38 | (Wang *et al.* 2019) |
| *G. hirsutum* TM1 (HAU v1) | 97.70% | 14.50% | 83.20% | 0.50% | 1.80% | 10.61 | (Wang *et al.* 2019) |

Genome annotation produced 40,181 transcripts representing 38,378 unique genes. Comparatively, the reference sequences for the related diploids *G. raimondii* (Paterson *et al.* 2012) and *G. arboreum (*Du *et al. 2018)* recovered 37,223 and 40,960 genes, respectively. Ortholog analysis between *G. longicalyx* and both diploids suggests a simple 1:1 relationship between a single *G. longicalyx* gene and a single *G. raimondii* or *G. arboreum* gene for 67-68% of the *G. longicalyx* genes (25,637 and 26,249 genes, respectively; Table 3). Approximately 3-4% of the *G. longicalyx* genome (i.e., 1,153-1,438 genes) are in “one/many” (Table 3) relationships whereby one or more *G. longicalyx* gene model(s) matches one or more *G. raimondii* or *G. arboreum* gene model(s). The remaining 5,009 genes were not placed in orthogroups with any other cotton genome, slightly higher than the 2,016 - 2,556 unplaced genes in the other diploid species used here. While this could be partly due to genome annotation differences in annotation pipelines, it is also likely due to differences in the amount of RNA-seq available for each genome.

**Table 3.** Orthogroups between *G. longicalyx* and two related diploid species. Numbers of genes are listed and percentages are in parentheses. Relationships listed in the last four lines of the table represent one/many G. longicalyx genes relative to one or many genes from G. arboreum or G. raimondii.

|  | *G. longicalyx* | *G. arboreum* | *G. raimondii* |
| --- | --- | --- | --- |
| Number of genes | 38,378 | 40,960 | 37,223 |
| Genes in orthogroups | 33,369<br>(86.9%) | 38,404<br>(93.8%) | 35,207<br>(94.6%) |
| Unassigned genes | 5,009 (13.1%) | 2,556 (6.2%) | 2,016 (5.4%) |
| Orthogroups containing species | 26,591<br>(78.5%) | 29,763<br>(87.8%) | 29,153<br>(86.0%) |
| Genes in species-specific orthogroups | 74 (0.2%) | 0 | 8 (0.0%) |
| 1-to-1 relationship |  | 26,249<br>(70.5%) | 25,637<br>(68.9%) |
| 1-to-many relationship |  | 1,207 (3.2%) | 1,153 (3.1%) |
| many-to-1 relationship |  | 1,438 (3.9%) | 1,172 (3.1%) |
| many-to-many relationship |  | 513 (1.4%) | 290 (0.8%) |

### Repeats

Transposable element (TE) content was predicted for the genome, both by *de novo* TE prediction (Bailly-Bechet *et al.* 2014; Smit *et al.* 2015) and repeat clustering (Novák *et al.* 2010). Between 44 - 50% of the *G. longicalyx* genome is inferred to be repetitive by RepeatMasker and RepeatExplorer, respectively. While estimates for TE categories (e.g., DNA, Ty3/*gypsy*, Ty1/*copia*, etc.) were reasonably consistent between the two methods (Table 4), RepeatExplorer recovered nearly 100 additional megabases of putative repetitive sequences, mostly in the categories of Ty3/*gypsy*, unspecified LTR elements, and unknown repetitive elements. Interestingly, RepeatMasker recovered a greater amount of sequence attributable to Ty1/*copia* and DNA elements (Table 4); however, this only accounted for 22 Mb (less than 20% of the total differences over all categories). The difference between methods with respect to each category and the total TE annotation is relatively small and may be attributable to a combination of methods (homology-based TE identification method versus similarity clustering), the under-exploration of the cotton TE population, and sensitivity differences in each method with respect to TE age/abundance.

**Table 4.** Comparison between repeat quantification methods for the *G. longicalyx* genome. Amounts are given in megabases (Mb).

|  | RepeatExplorer | RepeatMasker/OneCode |
| --- | --- | --- |
| LTR/Gypsy (Ty3) | 557 | 513 |
| LTR/Copia (Ty1) | 39 | 48 |
| LTR, unspecified | 44 | 0 |
| DNA (all element types) | 2.3 | 15 |
| unknown | 18 | 0 |
| Total repetitive clustered | 660 | 575 |

Because the RepeatExplorer pipeline allows simultaneous analysis of multiple samples (i.e., co-clustering), we used that repeat profile for both description and comparison to the closely related sister species, *G. herbaceum* and *G. arboreum* (from subgenus *Gossypium*). Relative to other cotton species, *G. longicalyx* has an intermediate amount of TEs, as expected from its intermediate genome size (1311 Mb; genome size range for *Gossypium* diploids = 841 - 2778 Mb). Approximately half of the genome (660 Mb) is composed of repetitive sequences, somewhat less than the closely related sister (A-genome) clade, which are slightly bigger in total size and have slightly more repetitive sequence (~60% repetitive; Table 5). Over 80% of the *G. longicalyx* repetitive fraction is composed of Ty3/*gypsy* elements, a similar proportion to the proportion of Ty3/*gypsy* in subgenus *Gossypium* genomes. Most other element categories were roughly similar in total amount and proportion between *G. longicalyx* and the two species from subgenus *Gossypium* (Figure 2).

**Table 5.**
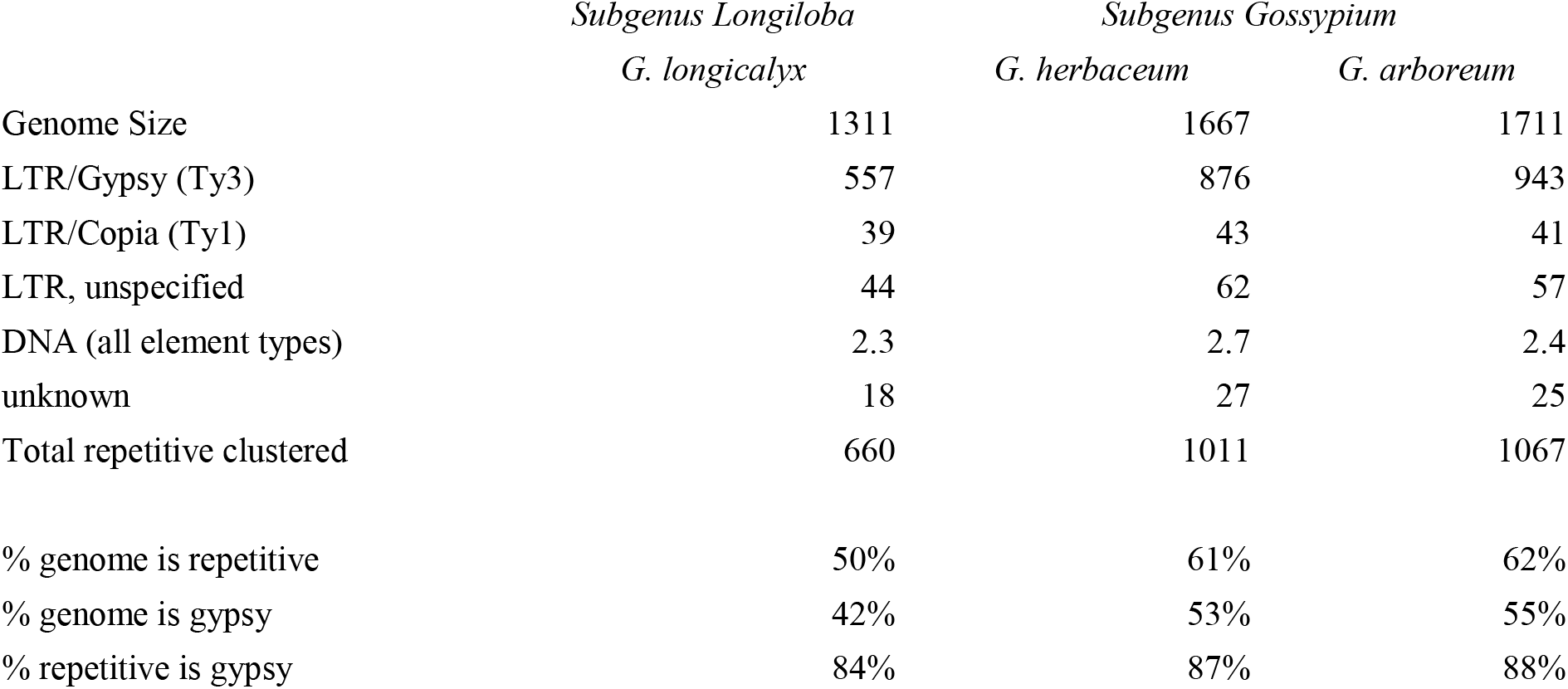
Transposable element content in *G. longicalyx* versus the sister clade (section *Gossypium*)

**Figure 2.**
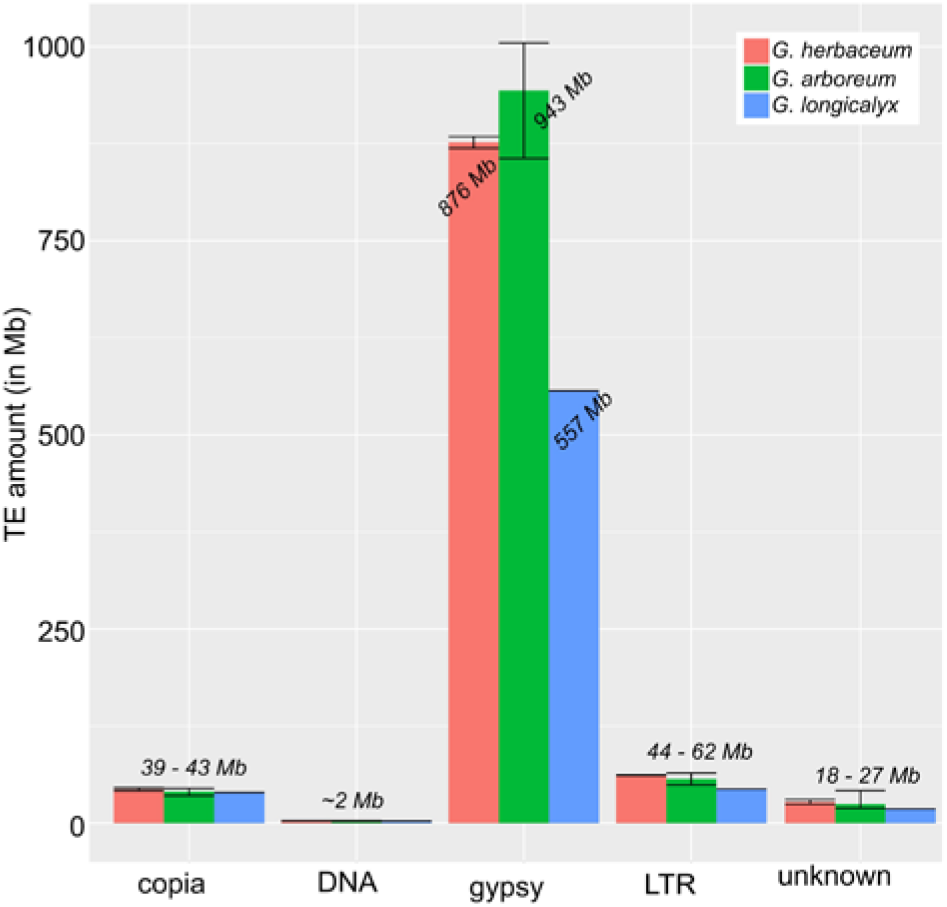
Repetitive content in *G. longicalyx* relative to the related diploid species *G. herbaceum* and *G. arboreum*.

### Chromatin accessibility in *G. longicalyx*

We performed ATAC-seq to map accessible chromatin regions (ACRs) in leaves. Two replicated ATAC-seq libraries were sequenced to ~25.7 and ~45.0 million reads per sample. The strand cross-correlation statistics supported the high quality of the ATAC-seq data, and the correlation of mapping read coverages (Pearson *r* = 0.99 and Spearman *r* = 0.77) suggested a high level of reproducibility between replicates (Supplemental Table 1). A total of 28,030 ACRs (6.4 Mb) were identified ranging mostly from 130 bp to 400 bp in length, which corresponds to ~0.5% of the assembled genome size (Supplemental Table 2). The enrichment of ACRs around gene transcription start sites (Supplemental Figure 1) suggested that these regions were functionally important and likely enriched with *cis*-regulatory elements. Based on proximity to their nearest annotated genes, these ACRs were categorized as genic (gACRs; overlapping a gene), proximal (pACRs; within 2 Kb of a gene) or distal (dACRs; >2 Kb from a gene). The gACRs and pACRs represented 12.2% and 13.2% of the total number of ACRs (952 Kb and 854 Kb in size, respectively), while approximately 75% (4.6 Mb) were categorized as dACRs, a majority of which were located over 30 Kb from the nearest gene (Figure 3). This high percentage of dACRs is greater than expected (~40% of 1 GB genome) given previous ATAC-seq studies in plants (Lu *et al.* 2019; Ricci *et al.* 2019) and may reflect challenges in annotating rare transcripts. While more thorough, species-specific RNA-seq will improve later annotation versions and refine our understanding of ACR proximity to genes, we do note that our observation of abundant dACRs and potentially long-range *cis*-regulatory elements is consistent with previous results (Lu *et al.* 2019; Ricci *et al.* 2019) The dACRs discovered here were the most GC-rich, followed by gACRs and pACRs (52%, 46%, and 44%, respectively), all of whom had GC contents significantly higher than randomly selected control regions with the same length distribution (Figure 3d). Because high GC content is associated with several distinct features that can affect the *cis*-regulatory potential of a sequence (Landolin *et al.* 2010; Wang *et al.* 2012), these results support the putative regulatory functions of ACRs.

**Figure 3.**
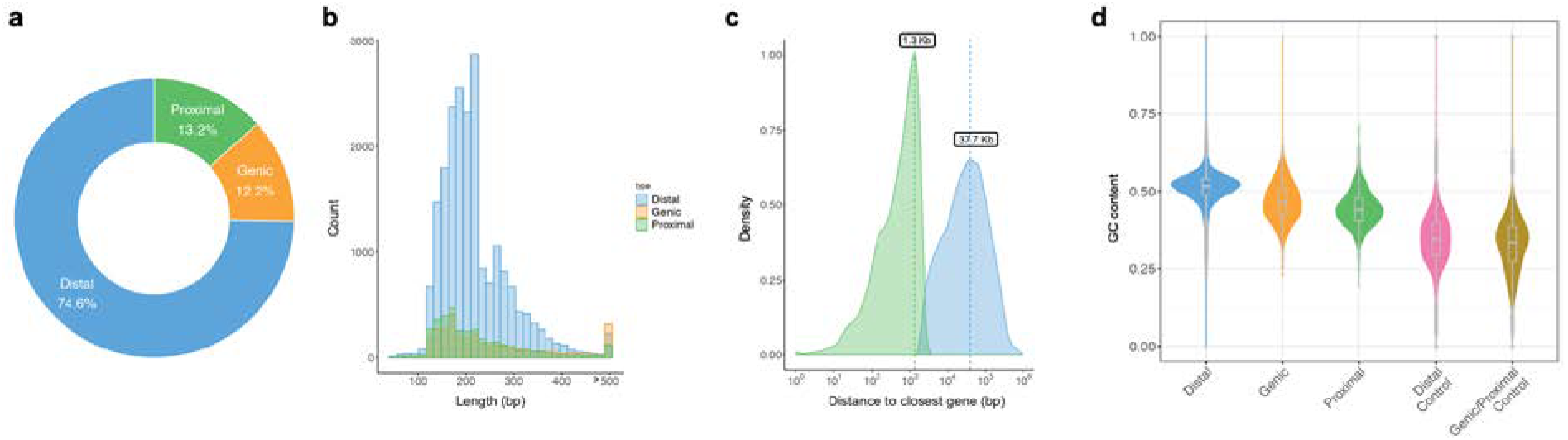
Accessible chromatin regions (ACRs) in the *G. longicalyx* genome. **a**. Categorization of ACRs in relation to nearest gene annotations - distal dACRs, proximal pACRs, and genic gACRs. **b**. Length distribution of ACRs that were identified by both HOMER and MACS2 contained within various genomic regions. **c**. Distance of gACRs and pACRs to nearest annotated genes. **d**. Boxplot of GC content in ACRs and control regions.

### Genomics of *G. longicalyx* reniform nematode resistance

Reniform nematode is an important cotton parasite that results in stunted growth, delayed flowering and/or fruiting, and a reduction in both yield quantity and quality (Robinson 2007; Khanal *et al.* 2018). While domesticated cotton varieties are largely vulnerable to reniform nematode (Robinson *et al.* 1997), nematode resistance is found in some wild relatives of domesticated cotton, including *G. longicalyx*, which is nearly immune (Yik and Birchfield 1984). Recent efforts to elucidate the genetic underpinnings of this resistance in *G. longicalyx* (i.e., Ren^Lon^) identified a marker (BNL1231) that consistently cosegrates with resistance and is flanked by the SNP markers Gl_168758 and Gl_072641 (Dighe *et al.* 2009; Zheng *et al.* 2016). Located in chromosome 11, this region contains one or more closely-linked nearly dominant gene(s) (Dighe *et al.* 2009) that confer hypersensitivity to reniform infection (Khanal *et al.* 2018), resulting in the “stunting” phenotype; however, the possible effects of co-inherited R-genes has not been eliminated. Because the introgressed segment recombines at reduced rates in interspecific crosses, it has been difficult to fine-map the gene(s) of interest. Additionally, progress from marker-assisted selection has been lacking, as no recombinants have possessed the desired combination of reniform resistance and “non-stunting” (Zheng *et al.* 2016). Therefore, more refined knowledge of the position, identity of the resistance gene(s), mode(s) of immunity and possible causes of “stunting” will likely catalyze progress on nematode resistance.

BLAST analysis of the three Ren^Lon^-associated markers (above) to the assembled *G. longicalyx* genome identifies an 850 kb region on chromosome F11 (positions 94747040..95596585; Figure 4) containing 52 predicted genes (Supplemental Table 3). Functional annotation reveals that over half of the genes (29, or 56%) are annotated as “TMV resistance protein N-like” or similar. In tobacco, TMV resistance protein N confers a hypersensitive response to the presence of the tobacco mosaic virus (TMV; (Erickson *et al.* 1999). Homologs of this gene in different species can confer resistance to myriad other parasites and pathogens, including aphid and nematode resistance in tomato (Rossi *et al.* 1998); fungal resistance in potato (Hehl *et al.* 1999) and flax (Ellis *et al.* 2007); and viral resistance in pepper (Guo *et al.* 2017). Also included in this region are 6 genes annotated as strictosidine synthase-like (SSL), which may also function in immunity and defense (Sohani *et al.* 2009). While the six SSL-like genes are tandemly arrayed without disruption, several other genes are intercalated within the array of TMV resistance-like genes, including the 6 SSL-like genes (Supplemental Table 3).

**Figure 4.**
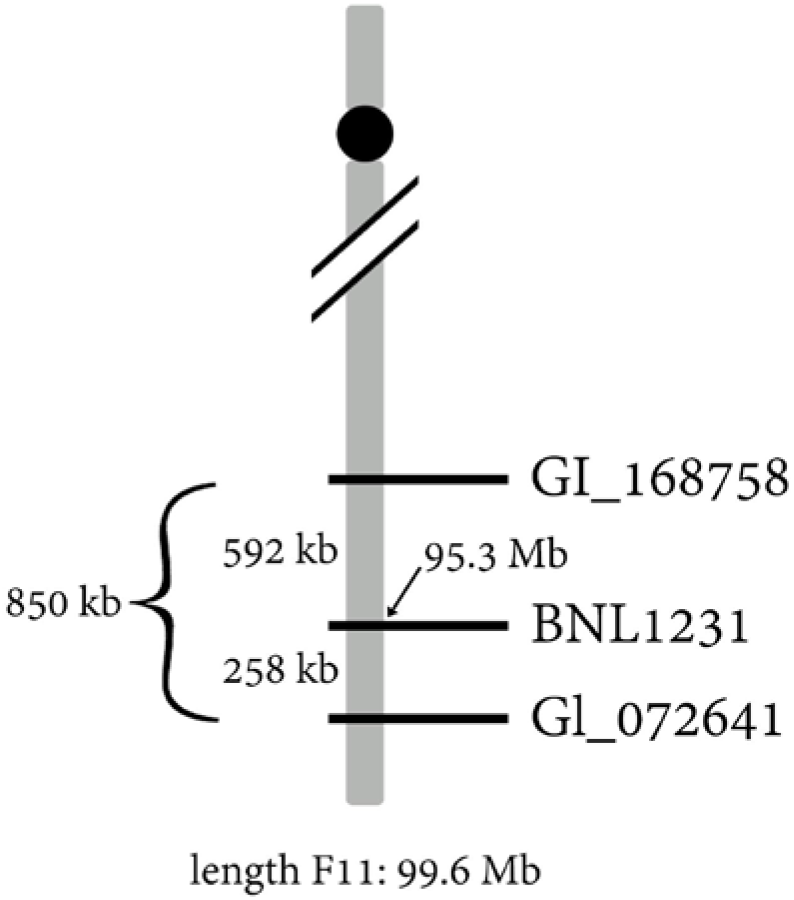
Diagram of the Ren^Lon^ region in *G. longicalyx*. Marker BNL1231, which co-segregates with nematode resistance, is located at approximately 95.3 Mb on chromosome F11.

Because there is agronomic interest in transferring nematode resistance from *G. longicalyx* to other species, we generated orthogroups between *G. longicalyx*, the two domesticated polyploid species (i.e., *G. hirsutum* and *G. barbadense*), and their model diploid progenitors (*G. raimondii* and *G. arboreum*; Supplemental Table 4; Supplemental File 2). Interestingly, many of the defense-relevant *G. longicalyx* genes in the Ren^Lon^ region did not cluster into orthogroups with any other species (15 out of 38; Table 6), including 11 of the 29 TMV resistance-related genes in the Ren^Lon^ region, and fewer were found in syntenic positions in *G. arboreum*. Most of the TMV resistance-related genes that cluster between *G. longicalyx* and other *Gossypium* species are present in a single, large orthogroup (OG0000022; Table 4), whereas the remaining TMV-resistance like genes from *G. longicalyx* are commonly in single gene orthogroups. Since disease resistance (R) proteins operate by detecting specific molecules elicited by the pathogen during infection (Martin *et al.* 2003), the increased copy number and variability among the *G. longicalyx* TMV-resistance-like genes may suggest specialization among copies.

**Table 6:**
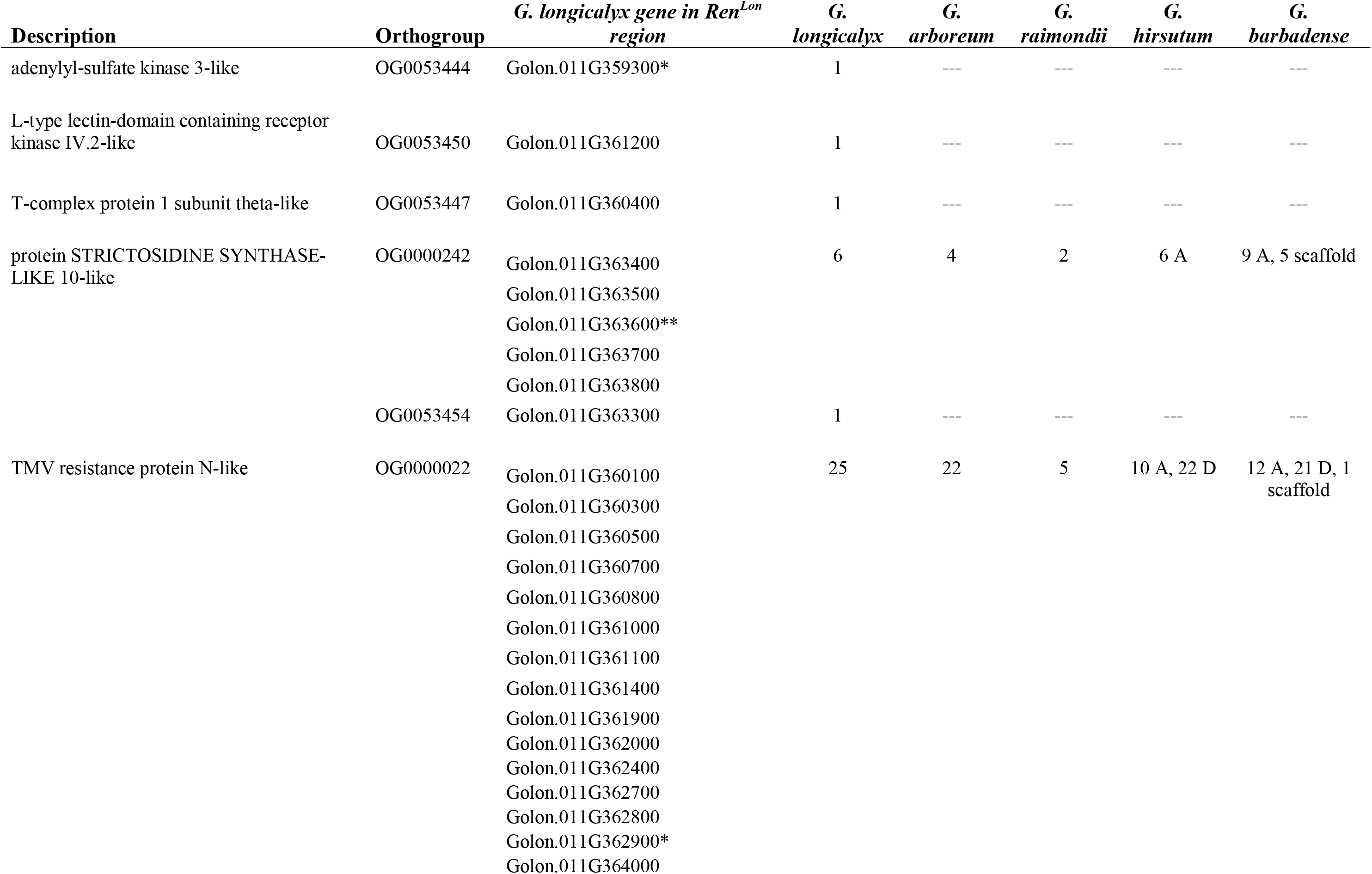
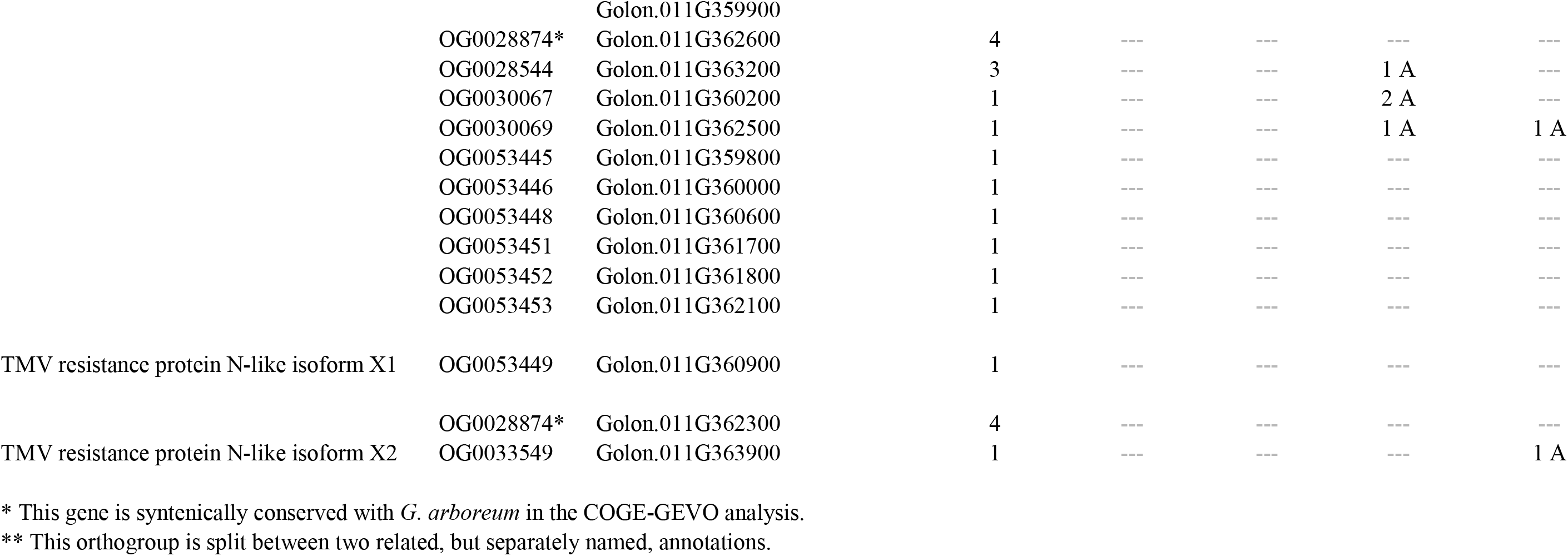
Orthogroup identity (by Orthofinder) for defense-related genes in the Ren^Lon^ region and the copy number per species. In *G. longicalyx*, this number includes genes found outside of the Ren^Lon^ region. *G. hirsutum* and *G. barbadense* copy numbers are split genes found on the A or D chromosomes, or on scaffolds/contigs not placed on a chromosome.

### Comparative genomics and the evolution of spinnable fiber

Cotton fiber morphology changed dramatically between *G. longicalyx* and its sister clade, composed of the A-genome cottons *G. arboreum* and *G. herbaceum*. Whereas *G. longicalyx* fibers are short and tightly adherent to the seed, A-genome fibers are longer and suitable for spinning. Accordingly, there has been interest in the changes in the A-genome lineage that have led to spinnable fiber (Hovav *et al.* 2008; Paterson *et al.* 2012). Progress here has been limited by the available resources for *G. longicalyx*, relying on introgressive breeding (Nacoulima *et al.* 2012), microarray expression characterization (Hovav *et al.* 2008), and SNP-based surveys (Paterson *et al.* 2012) of *G. longicalyx* genes relative to *G. herbaceum*. As genomic resources and surveys for selection are becoming broadly available for the A-genome cottons, our understanding of the evolution of spinnable fiber becomes more tangible by the inclusion of *G. longicalyx*.

Whole-genome alignment between *G. longicalyx* and the closely related *G. arboreum* (domesticated for long fiber) shows high levels of synteny and overall sequence identity (Figure 5). In general, these two genomes are largely collinear, save for scattered rearrangements and several involving chromosomes 1 and 2; these latter may represent a combination of chromosomal evolution and/or misassembly in one or both genomes. Notably, comparison of *G. longicalyx* to other recently published genomes (Supplemental Figures 2-5) suggests that an inversion in the middle of *G. longicalyx* Chr01 exists relative to representatives of the rest of the genus; however, the other structural rearrangements are restricted to *G. arboreum* and its derived A subgenome in *G. hirsutum* and *G. barbadense*, suggesting that these differences are limited to comparisons between *G. longicalyx* and A-(sub)genomes.

**Figure 5:**
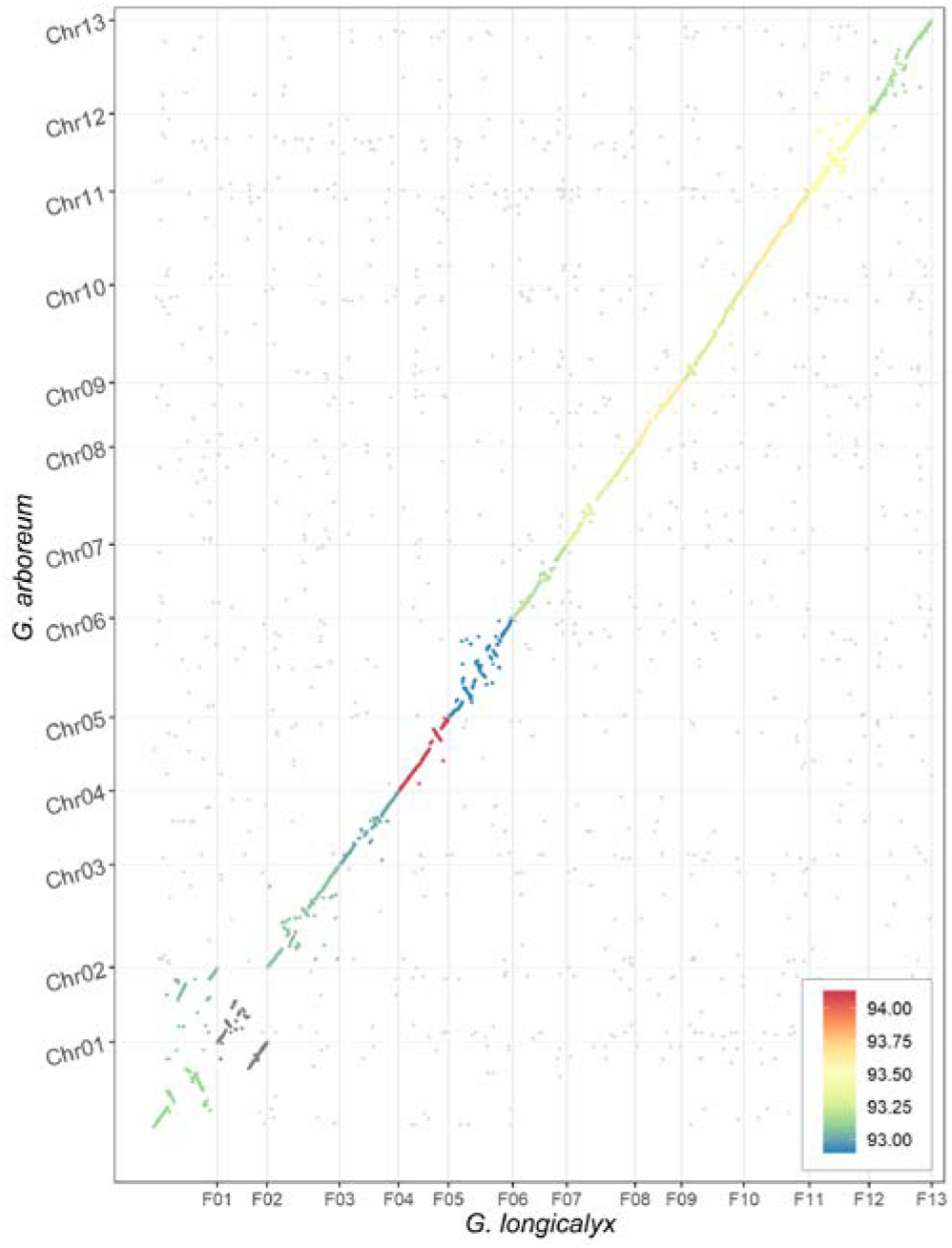
Synteny between *G. longicalyx* and domesticated *G. arboreum*. Mean percent identity is illustrated by the color (93-94% identity from blue to red), including intergenic regions.

Genic comparisons between *G. longicalyx* and *G. arboreum* suggests a high level of conservation. Orthogroup analysis finds a one-to-one relationship between these two species for over 70% of genes. Most of these putative orthologs exhibit <5% divergence (p-distance) in the coding regions, with over 50% of all putative orthologs exhibiting less than 1.5% divergence. Comparatively, the median divergence for putative orthologs between *G. longicalyx* and the more distantly related *G. raimondii* is approximately 2%, with ortholog divergence generally being higher in the *G. raimondii* comparison (Figure 6).

**Figure 6:**
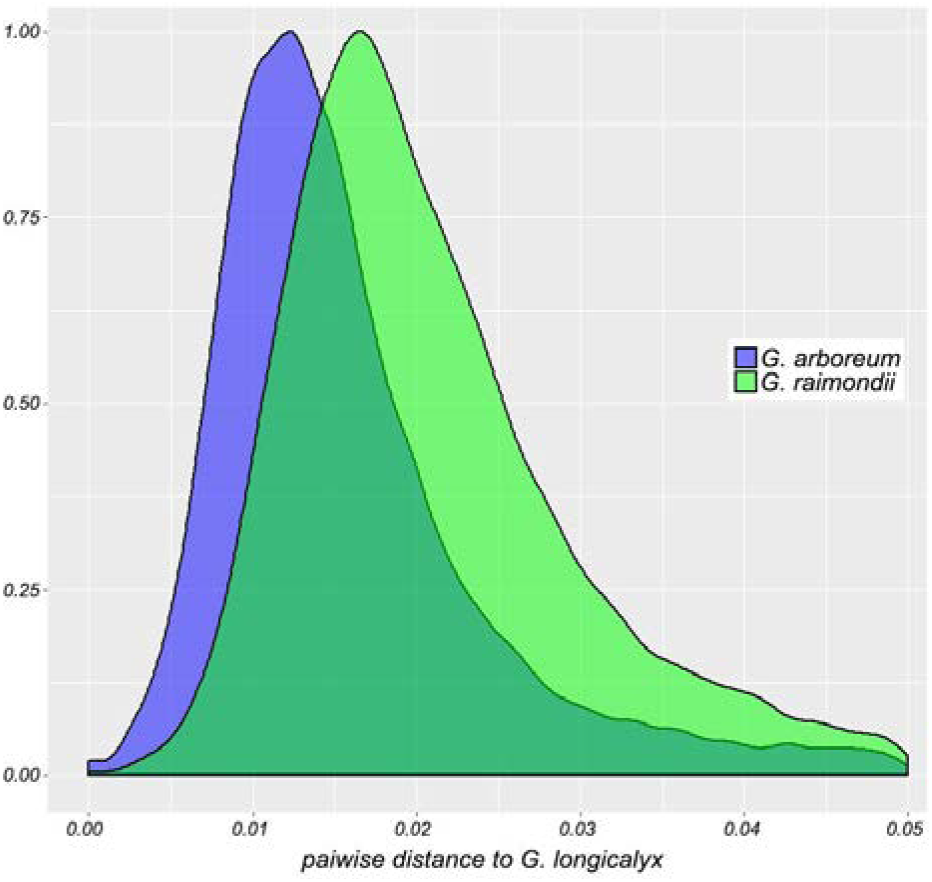
Distribution of pairwise p-distances between coding regions of predicted orthologs (i.e., exons only, start to stop) between *G. longicalyx* and either *G. arboreum* (blue) or *G. raimondii* (green). Only orthologs with <5% divergence are shown, which comprises most orthologs in each comparison.

Because *G. longicalyx* represents the ancestor to spinnable fiber, orthogroups containing only *G. arboreum* or polyploid A-genome gene annotations may represent genes important in fiber evolution. Accordingly, we extracted 705 *G. arboreum* genes from orthogroups composed solely of *G. arboreum* or polyploid (*i.e., G. hirsutum* or *G. barbadense*) A-genome gene annotations for BLAST and functional annotation. Of these 705 genes, only 20 represent genes known to influence fiber, i.e., ethylene responsive genes (10), auxin responsive genes (5), and peroxidase-related genes (5 genes; Supplemental Table 4). While other genes on this list may also influence the evolution of spinnable fiber, identifying other candidates will require further study involving comparative coexpression network analysis or explicit functional studies.

## Conclusion

While several high-quality genome sequences are available for both wild and domesticated cotton species, each new species provides additional resources to improve both our understanding of evolution and our ability to manipulate traits within various species. In this report, we present the first *de novo* genome sequence for *G. longicalyx*, a relative of cultivated cotton. This genome not only represents the ancestor to spinnable fiber, but also contains the agronomically desirable trait of reniform nematode immunity. This resource forms a new foundation for understanding the source and mode of action that provides *G. longicalyx* with this valuable trait, and will facilitate efforts in understanding and exploiting it in modern crop species.

## Acknowledgements

We thank Emma Miller and Evan Long for technical assistance. We thank the National Science Foundation Plant Genome Research Program (Grant #1339412) and Cotton Inc. for their financial support. This research was funded, in part, through USDA ARS Agreements 58-6066-6-046 and 58-6066-6-059. Support for R.J.S and Z.L. was provided by NSF IOS-1856627 and the Pew Charitable Trusts. We thank BYU Fulton SuperComputer lab for their resources and generous support. We also thank ResearchIT for computational support at Iowa State University. We thank Rise Services for office accommodations in Orem, UT.

